# dms2dfe: Comprehensive Workflow for Analysis of Deep Mutational Scanning Data

**DOI:** 10.1101/072645

**Authors:** Rohan Dandage, Kausik Chakraborty

**Affiliations:** CSIR Institute of Genomics and Integrative Biology, Mathura Road Campus, New Delhi 110020, India.; Academy of Scientific and Innovative Research (AcSIR), New Delhi 110001, India.

## Abstract

High throughput genotype to phenotype (G2P) data is increasingly being generated by widely applicable Deep Mutational Scanning (DMS) method. dms2dfe is a comprehensive end-to-end workflow that addresses critical issue with noise reduction and offers variety of crucial downstream analyses. Noise reduction is carried out by normalizing counts of mutants by depth of sequencing and subsequent dispersion shrinkage at the level of calculation of preferential enrichments. In downstream analyses, dms2dfe workflow provides identification of relative selection pressures, potential molecular constraints and generation of data-rich visualizations.

**Availability:** dms2dfe is implemented as a python package and it is available at https://kc-lab.github.io/dms2dfe.

**Supplementary information:** Supplementary data are available at Bioinformatics online.

## 1 Introduction

Recent developments in high throughput mutagenesis and massively parallel DNA sequencing culminated in a method popularly known as Deep Mutational Scanning (DMS) (1,2). The broad appeal of DMS method is evident by the wide range of applicability (3) relevant to structural modeling (4), protein stability (5), substrate specificity (6–8), protein-protein interactions (9–14), mutational effects on organismal fitness (15–17), environmental effects (18), biology of viruses (19–21), fitness landscapes of antibiotic resistance (7,22) and proto-oncogenes(23). The distribution of fitness scores along fitness axis of fitness landscape i.e. distribution of fitness effects (DFE) is known as a powerful estimator of the underlying evolutionary dynamics and has been traditionally used to contextualize molecular evolution (24,25). One can characterize the nature of selection pressure in effect and potential susceptibility, evolvability or robustness of the individuals in the population by comparing DFEs.

Accounting and reducing for the noise is a major primary challenge in the analysis of differential count data. Adopting an alternative approach to the analysis than available tools (26–28), dms2dfe implements noise reduction utilizing both sequencing depth (Fig S1 A) and empirical Bayes shrinkage (29) (Fig S1 B). Along with upstream analysis to obtain variant counts, preferential enrichments and fitness estimates, dms2dfe offers utilities for downstream analysis such as identification of relative selection pressure, molecular constraints and visualizations of the data; thus constituting an end-to-end workflow. Comprehensively documented python programming library (API) (available at http://kc-lab.github.io/dms2dfe) provides a platform for further developments to adapt with advances and modifications in DMS experiments.

## 2 dms2dfe workflow

In a conventional DMS experiment, a pool of mutants is selected in a co-culture competition assay under selection pressure of interest and frequencies of the mutants are compared with a pool or mutants used as a reference (also referred as input or background). dms2dfe is applicable to all such approaches wherein a basic DMS experiment involves a comparative analysis of selected and reference pool of mutants. In addition to the shot gun and full length ultra-deep sequencing, dms2dfe also supports concatamer based approach (7) as well as multiplexing strategy using barcoded amplicons (30). Sequencing data whether aligned (.sam, .bam) or not (.fastq) can be provided as input of dms2dfe workflow. Alternatively, mutational data (frequencies, preferential enrichments or fitness score) can be provided to exclusively carry out downstream analyses.

As shown in Fig 1 A, in the comprehensive workflow of dms2dfe, following key aspects of the analysis of DMS experiment are addressed.

**Fig. 1.**
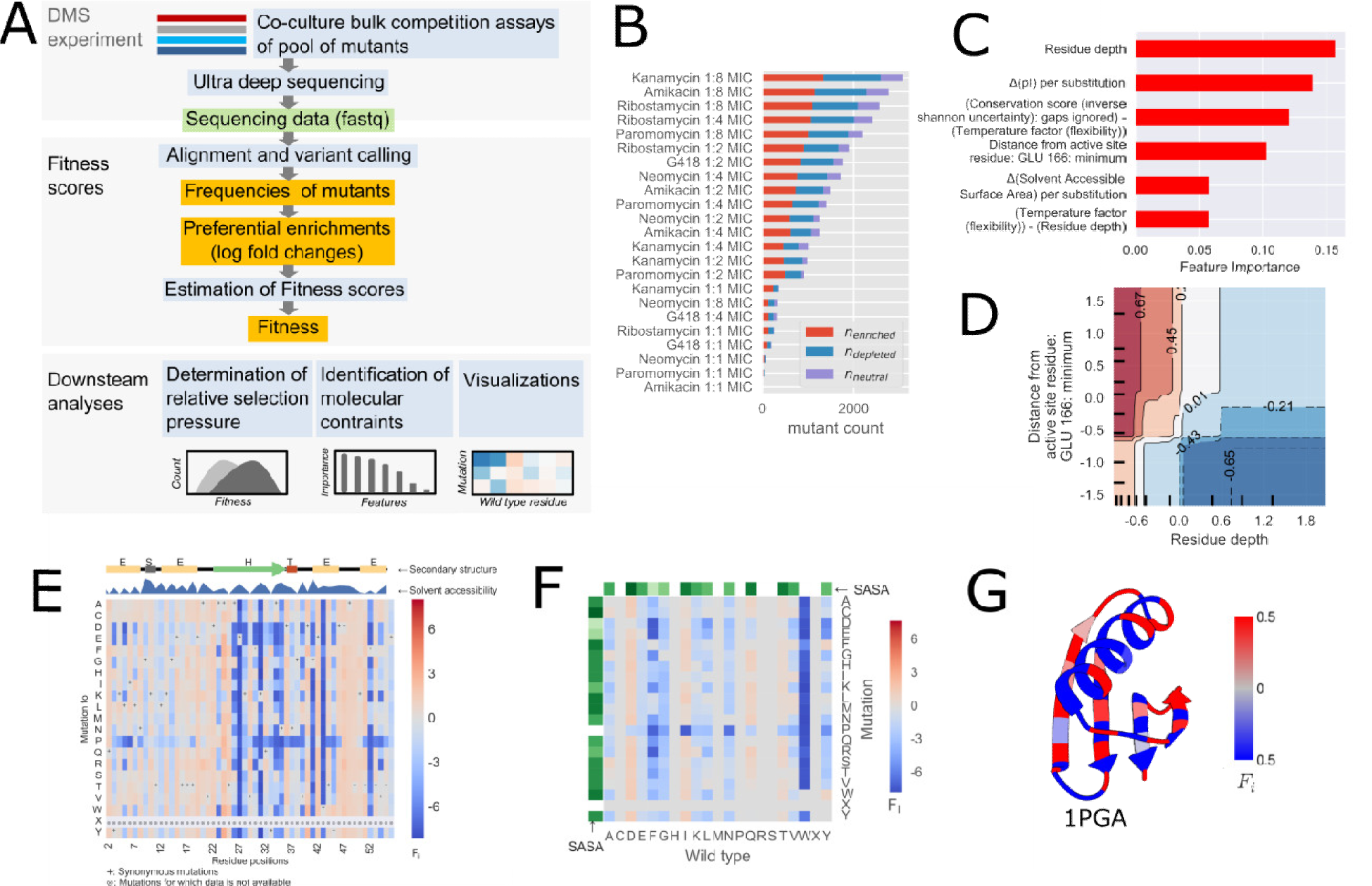
Workflow and functionalities of dms2dfe. (A) Sequencing data produced by DMS experiment is preprocessed, aligned and variants are identified. Preferential enrichments (log fold changes) are used to estimate Fitness scores according to several methods (see S1 *Supporting methods*). Among downstream analyses, dms2dfe provides methods to identify the selection pressure among multiple DFEs, determine molecular constraints of the fitness landscape and generate various visualizations of the mutational data. (B) Demonstration of determination of relative selection pressure in the form of counts of mutants classified as enriched (n_enriched_), depleted (n_depleted_), and neutral (n_neutral_). (C and D) Identification of molecular constraints of fitness data. (C) Feature importance of top features with highest feature importance as determined a binary classification model (S1 Materials and Methods) are dependence of the predictors is scaled along z axis. Axis ticks in represent decile values of the features. (E-F) Visualizations of mutational data in three different formats i.e. (E) fitness per mutations (F) average fitness per substitutions and (G) average fitness per positions of the protein. In panel E, each locus of the heatmap represents fitness of individual mutant. In panel F, each locus of the heatmap represents average fitness score per substitution. In panel G, average fitness score per residue position is projected onto PDB structure (1PGA) of the protein. Here, for generating plot B, APH2 dataset was used. For generating plots C and D, APH2 dataset was used. For generating plot E-H, GB1 dataset was used.

### 2.1 Identification of relative selection pressure

As previously implemented (8), we classify the statistically neutral (least effect) mutants upper and lower thresholds from the preferential enrichments of reference (unselected) pool. Depending on the nature of experiment, fitness scores can be estimated by one of several methods based on preferential enrichments of wild type alleles or otherwise preferential enrichments themselves can be used as fitness scores (see S1 *Supporting Methods*). So, mutants can be classified as beneficial, deleterious or neutral as follows,

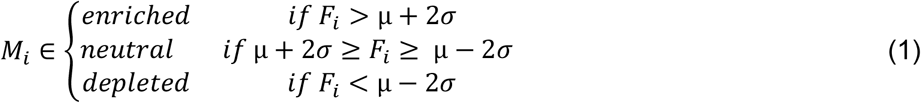

where, *M_i_* is i^th^ kind of survived mutant and *F_i_* is its respective fitness score. Note that class neutral exclusively represents the synonymous mutants. Metrics such as relative number of mutations under the classes of fitness (Fig 1B), relative survivability i.e. difference in number of mutants survived in test condition with respect to control condition (Δn) and relative fitness changes i.e. difference in average fitness of mutants in test condition with respect to control condition (ΔF) assist users to identify the directionality and strength of the relative selection pressures.

### 2.2 Identification of molecular constraints

Due to multiple reasons (31), effects of mutation at one site are dependent on residues at other sites. This results in a highly interdependent ensemble of molecular constraints which is difficult to decipher (32). Here we use ensemble machine learning algorithm - Gradient Boosting which allows modeling of the complex mutational data. It provides critical advantage of determination by relative importances of the features (Fig 1 C) and partial dependence i.e. marginal effect of features on the target i.e. fitness scores (Fig S2). Such information would help users in contextualizing G2P interactions in terms of how fitness scores are mechanistically constrained by molecular constraints (Fig S1 D).

### 2.3 Visualizations

Fitness data too can be visualized in three different formats. Fitness per individual mutations is visualized in the form of a heatmap also called sequence-function map, wherein each locus represents fitness of individual mutant (Fig 1 E). Fitness per substitutions too can be visualized in the form of matrix of average fitness scores per individual substitution (Fig 1 F). Average fitness values per position are projected onto the PDB structure by utilizing visualization modules of UCSF-Chimera (34)(Fig 1 G). Alongside, an integrated visualization of the change in frequencies of mutants at the level of individual mutations, substitutions and positions is generated (Fig S1 B).

## 3 Conclusions

dms2dfe is a tunable, open-source workflow that integrates state-of-the-art methods of genomics for analysis of DMS data. Accounting variable sequencing depth across the length of the gene and implementing dispersion shrinkage though empirical Bayes approach results in reduction in the noise associated with empirical data. Downstream analyses such as identification of relative selection pressures, identification of molecular constraints, and intuitive visualizations help in contextualizing the DMS data. Collectively, dms2dfe offers experimenters a comprehensive solution for the analysis of DMS data and an open-source platform for future developments to accommodate potential modifications of DMS method.

## Additional files

S1 text: Supporting Methods, Figures and Tables.

## Authors’ contributions

RD and KC designed the method. RD implemented the method. RD and KC wrote the manuscript. All authors read and approved the final manuscript.

## Acknowledgments

We acknowledge CSIR for its funding through EMPOWER project and infrastructural support from CSIR IGIB. R.D. acknowledges UGC for graduate funding. We thank the members of KC lab for critically reviewing the manuscript.

